# Recombination of 2Fe-2S ferredoxins reveals differences in the inheritance of thermostability and midpoint potential

**DOI:** 10.1101/2020.05.29.122317

**Authors:** Ian J. Campbell, Dimithree Kahanda, Joshua T. Atkinson, Othneil N. Sparks, Jinyoung Kim, Chia-Ping Tseng, Rafael Verduzco, George N. Bennett, Jonathan J. Silberg

## Abstract

Homologous recombination can be used to create enzymes that exhibit distinct activities and stabilities from proteins in nature, allowing researchers to overcome component limitations in synthetic biology. To investigate how recombination affects the physical properties of an oxidoreductase that transfers electrons, we created ferredoxin (Fd) chimeras by recombining distantly-related cyanobacterial and cyanomyophage Fds that present similar midpoint potentials but distinct thermostabilities. Fd chimeras having a wide range of amino acid substitutions retained the ability to coordinate an iron-sulfur cluster, although their thermostabilities varied with the fraction of residues inherited from each parent. The midpoint potentials of chimeric Fds also varied. However, all of the synthetic Fds exhibited midpoint potentials outside of the parental protein range. Each of the chimeric Fds could also be used to build synthetic pathways that support electron transfer between Fd-NADP reductase and sulfite reductase in *Escherichia coli*, although the chimeric Fds varied in the expression required to support similar levels of cellular electron transfer. These results show how recombination can be used to rapidly diversify the physical properties of protein electron carriers and reveal differences in the inheritance of thermostability and electrochemical properties. Furthermore, they illustrate how electron transfer efficiencies of chimeric Fds can be rapidly evaluated using a synthetic electron transfer pathway.

## INTRODUCTION

Ferredoxins (Fds) are small soluble protein electron carriers that evolved to shuttle electrons in organisms across the tree of life, with some cells having genomes that encode as many as three dozen paralogs^1^. Fds transfer electrons between a wide range of partner oxidoreductases, ranging from proteins involved in light harvesting and nutrient assimilation to steroid synthesis and porphyrin metabolism^2–7^. Among all protein electron carriers, Fds present some of the lowest midpoint reduction potentials (E°), with [2Fe-2S] Fds ranging from −150 to −500 mV and [4Fe-4S] Fds ranging from −200 to −650 mV^2^. While Fds are thought to control cellular electron transfer (ET) between their diverse partners by evolving sequences with distinct partner affinities and electrochemical properties, we cannot yet anticipate a priori how changes in Fd primary structure alter these biochemical and biophysical properties for synthetic biology applications.

Homologous recombination can be used in the laboratory to study how primary structure controls protein function^8,9^. This approach is appealing to use for protein design because amino acid substitutions created by recombination are less disruptive than those created randomly, since the sequence blocks being swapped have already been selected by evolution for compatibility within native structures^10^. Recombination has been applied to a variety of enzymes including metal-containing oxidoreductases, such as cytochromes P450, laccases, and hydrogenases^11–14^. These studies have revealed that recombination can lead to innovation in metalloproteins, creating chimeras with distinct properties from the parent proteins being bred, including higher catalytic activity, distinct substrate specificity profiles, and altered stabilities^11–14^.

Only a small number of studies have examined the effects of recombination on Fd electron carriers. These efforts have largely focused on recombining Fds that are encoded by the same genome and exhibit high sequence identity. Amino acids substitutions created by recombination have yielded Fd chimeras with a range of properties, including decreased stability, solubility, and ET efficiencies with partner proteins^15–19^. In a few cases, recombination has yielded Fds with electrochemical properties that differ from the parental proteins^15–17^. In these studies, Fds with distinct E° were recombined, and the E° of the resulting chimeras were within the range of values bounded by the parental proteins, with the E° values depending upon the relative degree of structural similarity to each parent. This trend is similar to that observed when homologous proteins with distinct thermostabilities are recombined^20^. The extent to which this trend applies to other Fd chimeras has not been explored.

Biophysical studies have shown that changes in the hydrogen bonding network surrounding the Fd iron-sulfur cluster can lead to E° changes^21–23^. Because the number of hydrogen bonds disrupted by recombination is inversely correlated with parental protein sequence identity^13^, we hypothesized that recombination of distantly-related Fds would alter E° more dramatically than observed in prior studies. To test this idea, we created a set of Fd chimeras by recombining homologs from the cyanobacterium *Mastigocladus laminosus* Fd (ml-Fd1) and a cyanophage PSSM-2 (pssm2-Fd). These Fds were chosen because they have near-identical electrochemical properties (E° ≅ −340 mV), while their melting temperatures (T_m_) differ by >40°C^24–26^. In each chimera created, we examined iron-sulfur cluster coordination, E°, thermostability, and ET in a synthetic pathway within cells. Surprisingly, all of the chimeras coordinated iron-sulfur clusters and supported cellular ET. However, chimeras presented varying stabilities, and many chimeras presented E° that were outside of the parental range.

## RESULTS AND DISCUSSION

### Chimera design

ml-Fd1 and pssm2-Fd were targeted for design because they exhibit similar structures (RMSD = 0.4 Å) but differ in sequence at almost half of their native positions (Figure 1). Many of the native sites that differ in sequence also vary in charge (n=16) and possess functional groups that can form hydrogen bonds (n=8). Even with these differences, the parental proteins contain similar total numbers of hydrogen bonds, with ml-Fd1 having 123 hydrogen bonds and pssm2-Fd having 133 hydrogen bonds^25,27^. Because of the differences in the distribution of residues within the primary structures of each parent, recombination has the potential to disrupt up to 51 total residue-residue contacts, including contacts that participate in hydrogen bonds (n=38) and salt bridges (n=4), molecular interactions that contribute to Fd electrochemical properties^21–23^.

**Figure 1.**
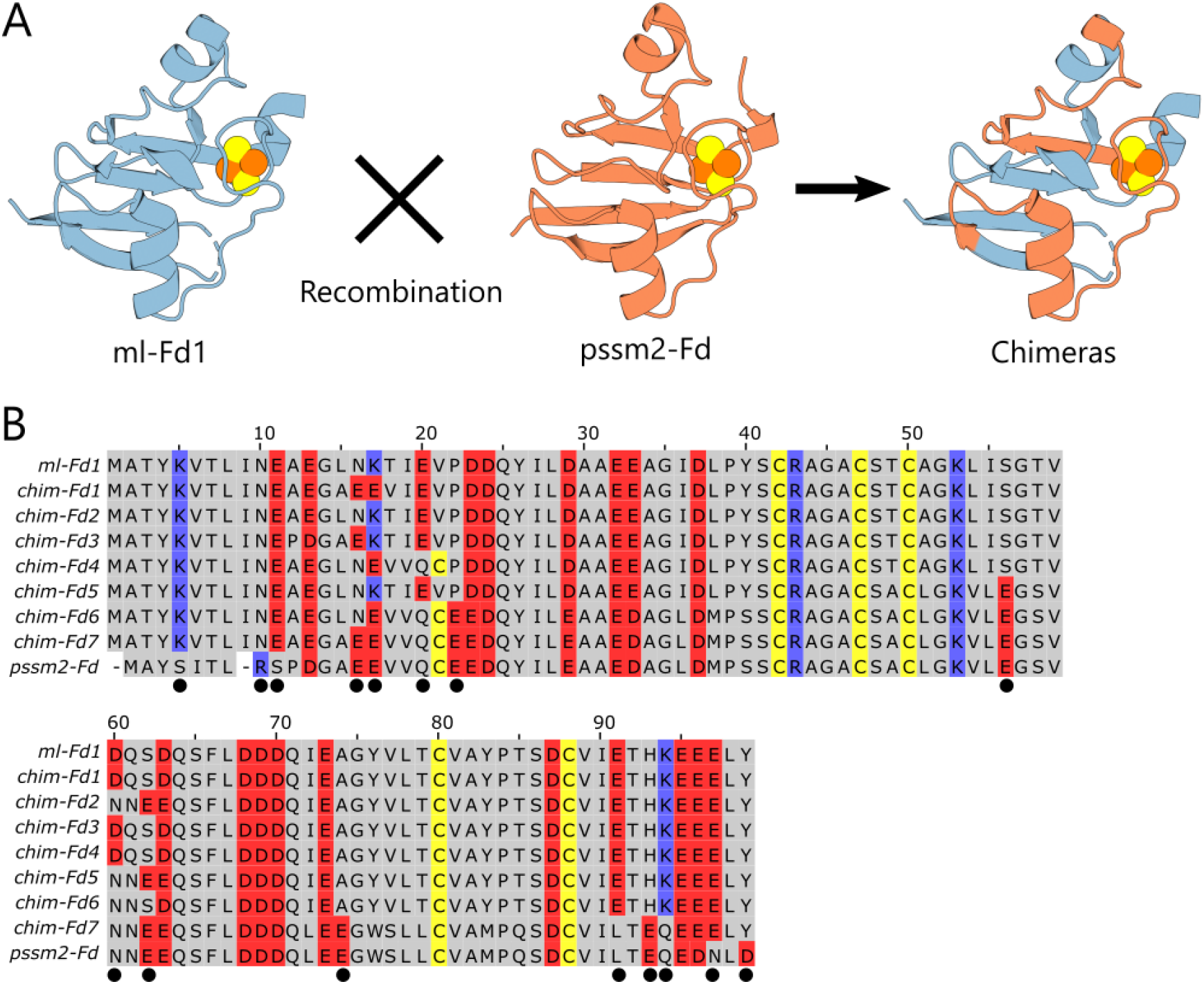
Comparison of parental and chimeric Fds. (**A**) Structures of ml-Fd1 (PDB: 1RFK) and pssm2-Fd (PDB: 6VJV) which were recombined to create chimeric Fds^25,27^. (**B**) Alignment of the chimeric Fds and the parental proteins reveal residues that vary in charge (black circles).

In total, we generated seven chimeric ferredoxins (chim-Fd) which are named numerically based on their mutational distance from ml-Fd1 (Table 1; Figure S1). A small set of chimeras was targeted to enable comprehensive in vitro characterization and allow for calibration of the disruptive nature of recombination as previously described^13,28^. Chimeras were created with a range of amino acid substitutions (4 to 34) relative to ml-Fd1. These chimeras exhibited small variation in their absolute numbers of positively and negatively charged residues such that they all maintain the characteristically low pI found in Fds^29–31^. However, the chimeras differed in the total number of disrupted residueresidue contacts (5 to 29), number of disrupted hydrogen bonds (5 to 23), and number of disrupted salt bridges (0 to 1).

**Table 1.** Physicochemical properties of Fd chimeras. Properties include isoionic point (pI), mutational distance from ml-Fd1 (m_ml-Fd1_), ratio of ellipticity at 427 to concentration (θ_427_/μM), midpoint reduction potential (E°), melting temperature (T_m_), and residue-residue contacts broken by recombination relative to most similar parent (C_broken_).

| Fd | $m_{\text{ml-Fd1}}$ | pI | $\theta_{427}/\mu\text{M}$ | $E^\circ$ (mV/SHE) | $T_m$ ($^\circ\text{C}$) | $C_{\text{broken}}$ |
| --- | --- | --- | --- | --- | --- | --- |
| ml-Fd1 | 0 | 3.9 | 0.28 | -344 | 72.6 | 0 |
| chim-Fd1 | 4 | 3.75 | 0.25 | -354 | 64.7 | 11 |
| chim-Fd2 | 4 | 3.94 | 0.21 | -330 | 66.5 | 5 |
| chim-Fd3 | 4 | 3.86 | 0.30 | -392 | 64.7 | 11 |
| chim-Fd4 | 5 | 3.79 | 0.23 | -396 | 67.3 | 17 |
| chim-Fd5 | 10 | 3.91 | 0.28 | -384 | 66.2 | 13 |
| chim-Fd6 | 20 | 3.81 | 0.27 | -360 | 42.6 | 29 |
| chim-Fd7 | 34 | 3.54 | 0.08 | -319 | 20 | 23 |
| pssm2-Fd | 47 | 3.5 | 0.25 | -336 | 28 | 0 |

### Cofactor content and thermostability

All of the chimeric Fds were purified using a combination of ion exchange and size exclusion chromatography. Each recombinant Fd presented a brown color immediately following purification, consistent with the presence of a 2Fe-2S cluster, although the final yields of each protein varying by >35-fold, with chim-Fd4 yielding 94 mg/L and chim-Fd7 yielding only 2.5 mg/L. To investigate which chimeric Fds contain iron-sulfur clusters, we measured their absorbance and circular dichroism (CD) spectra and compared them with spectra obtained using purified ml-Fd1 and pssm2-Fd. In all cases, the chim-Fds presented absorbance spectra with peaks (465, 420, and 330 nm) that are characteristic of [2Fe-2S]-bound Fds (Figure S2)^31–33^. Additionally the chim-Fds had CD spectra with ellipticity maxima (427 and 360 nm) and minima (505 and 560 nm) consistent with native holoFds (Figure 2)^32,33^. To compare chimera metallocluster content, we calculated the ratio of ellipticity at 427 nm to the concentrations determined using absorbance (Figure S3). While many of the chimeras presented ratios consistent with a high [2Fe-2S] occupancy (76 to 113% parental values), chim-Fd7 had a ratio corresponding to only 30% occupancy.

**Figure 2.**
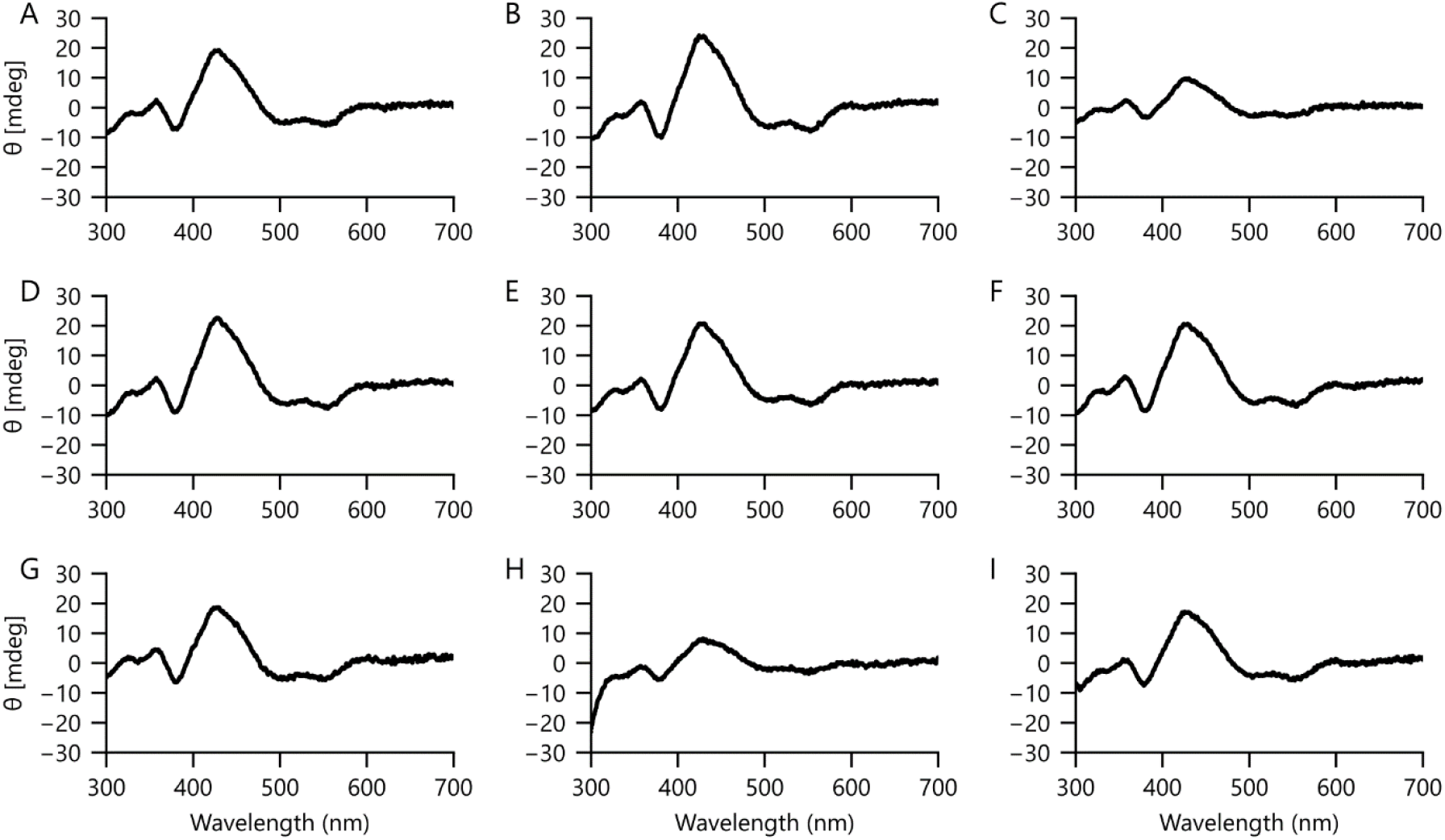
Circular dichroism spectra of chimeric Fds. The circular dichroism spectra of (**A**) ml-Fd1, (**B**) chim-Fd1, (**C**) chim-Fd2, (**D**) chim-Fd3, (**E**) chim-Fd4, (**F**) chim-Fd5, (**G**) chim-Fd6, (**H**) chim-Fd7, and (**I**) pssm2-Fd exhibits classic features of [2Fe-2S] Fds. Measurements were performed using 50 μM protein in TED buffer at 23°C.

The thermostabilities of parental Fds differ by 46°C, with pssm2-Fd and ml-Fd1 having T_m_ values of 28°C and 74°C, respectively^25,26^. To examine how the stabilities of the 2Fe-2S in the chim-Fds relate to the parental proteins, we measured how their ellipticities (427 nm) changed with increasing temperature (Figure 3). All of the chim-Fds exhibited temperature-dependent ellipticities. A majority of the chimeras exhibited midpoints for their loss of ellipticity at temperatures that are intermediate between the two parents, with the exception for chim-Fd7, which exhibited a lower midpoint value than pssm2-Fd. The five chimeras that are more closely related to ml-Fd1 (chim-Fd1, chimFd2, chim-Fd3, chim-Fd4, and chimFd-5) presented midpoint values in a narrow range (65 to 67°C) while the other chimeras exhibited lower values (43°C and 20°C).

**Figure 3.**
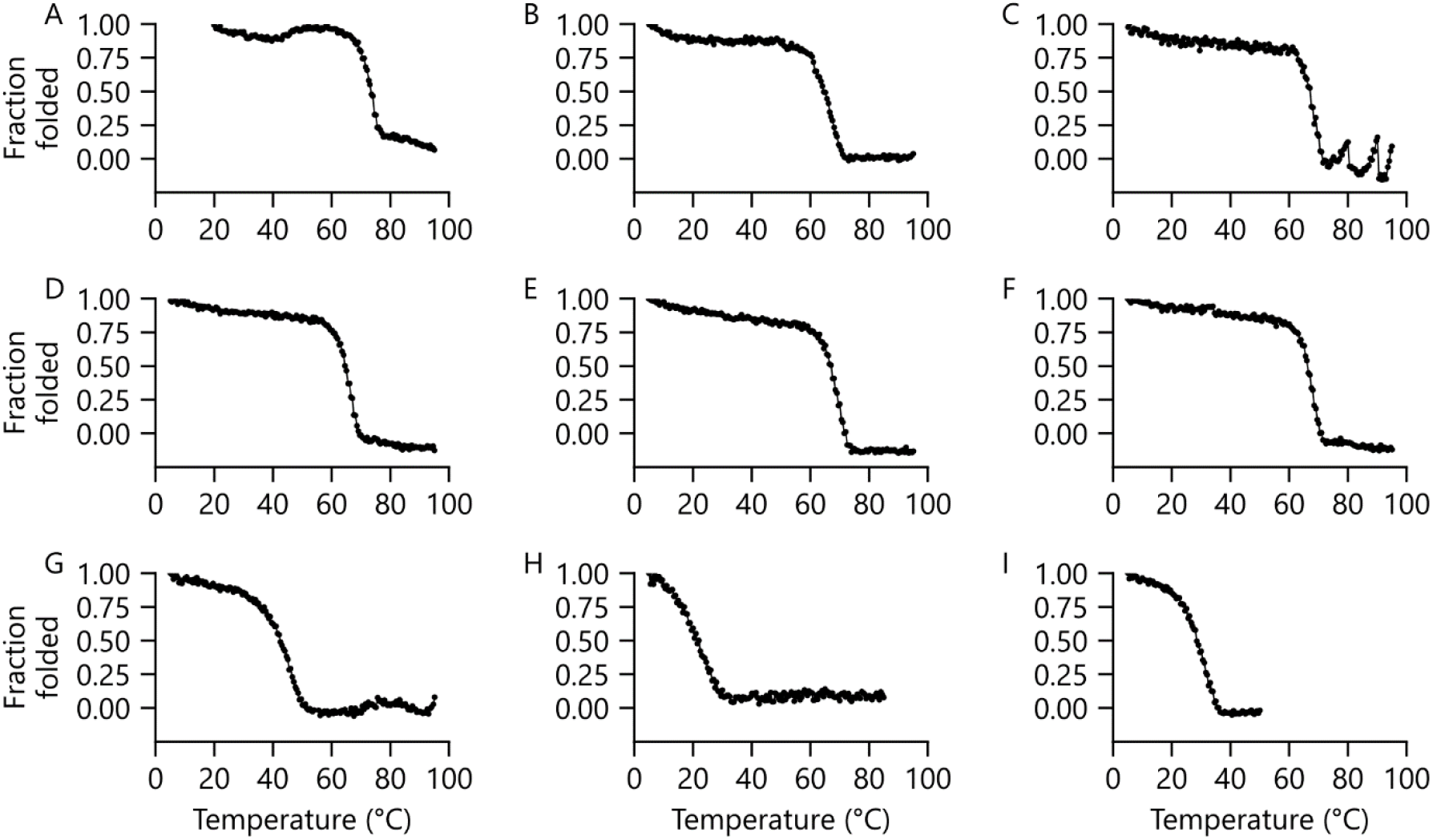
Effect of temperature on iron-sulfur cluster coordination. The thermal denaturation of (**A**) ml-Fd1, (**B**) chim-Fd1, (**C**) chim-Fd2, (**D**) chim-Fd3, (**E**) chim-Fd4, (**F**) chim-Fd5, (**G**) chim-Fd6, (**H**) chim-Fd7, and (**I**) pssm2-Fd. All experiments were performed by monitoring ellipticity (427 nm) of samples containing 50 μM protein in TED buffer at a scan rate of 1 C°/min.

### Electron transfer in a synthetic pathway

The parental Fds can both support ET from *Zea mays* FNR (zm-FNR) to *Zea mays* SIR (zm-SIR) within a synthetic cellular pathway^24,25,34^. In this cellular assay, ET is quantified by monitoring the growth of an *Escherichia coli* strain (EW11) that cannot grow on medium containing sulfate as the only sulfur source unless a Fd transfers electrons from FNR to SIR^34^. When a Fd transfers electrons from FNR to SIR, sulfite is reduced to sulfide to facilitate the synthesis of essential metabolites like cysteine and methionine. To test whether any of the chimeras support ET from FNR to SIR like the parents, we electroporated *E. coli* EW11 with a plasmid that expresses each chimeric Fd using an anhydrotetracycline (aTc) inducible promoter and a plasmid that constitutively expresses zm-FNR and zm-SIR. In all cases, cells expressing chimeric Fds showed significant growth following overnight incubations in the presence of aTc over the non-induced controls (Figure 4A). Five of the chimeras (chim-Fd2, -Fd3, -Fd4, -Fd5, and -Fd6) presented endpoint growth complementation that was similar to parental proteins. However, the growth enhancement with chim-Fd1 and chim-Fd7 were both lower than parental proteins, even though these chimeras are the two chim-Fds most closely related to the two different parental proteins.

**Figure 4.**
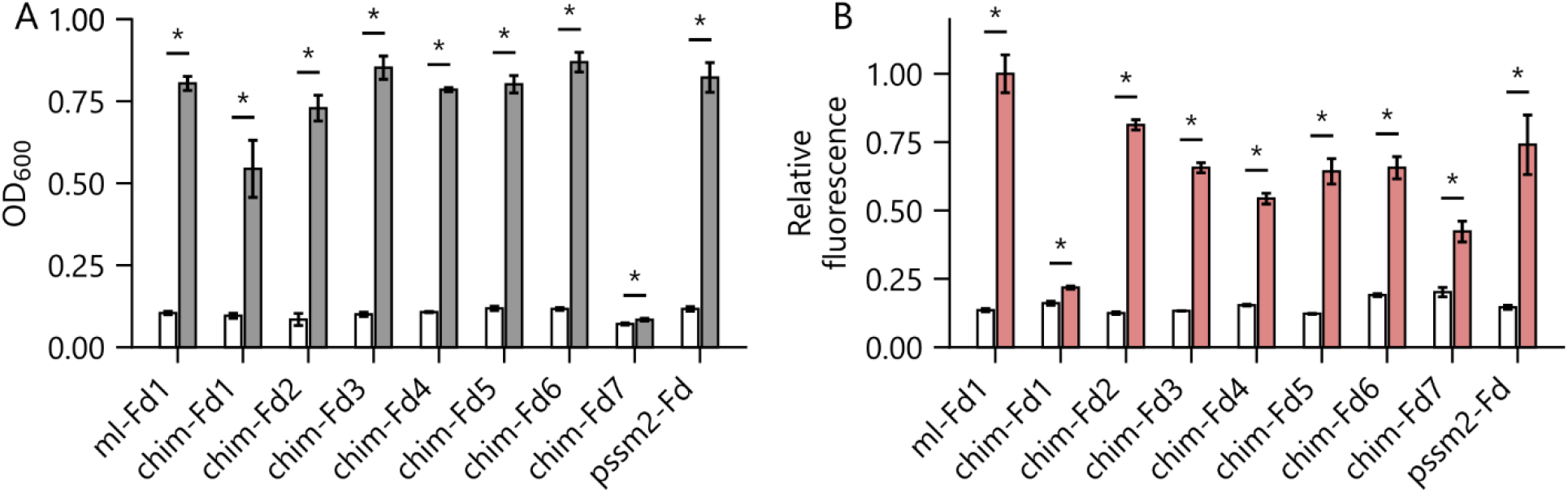
Chimeric Fds support ET between plant FNR and SIR. (**A**) Growth of *E. coli* EW11 in M9sa medium in the absence (white) and presence (gray) of aTc. Cells were cotransformed a vector that constitutively expresses zm-FNR and zm-SIR and different vectors that expresses each Fd using an aTc-inducible promoter. (**B**) *E. coli* EW11 was transformed with vectors for expressing Fd-RFP fusions, and whole cell fluorescence was measured in uninduced (white) and induced (red) cells. Relative fluorescence represents emission divided by OD_600_ and normalized to the maximum value observed. All experiments were performed in triplicate, and an independent two-tailed t-test was used to compare values ±aTc (α=0.05). Significant differences (p<0.05) are noted with asterisks.

Differences in chim-Fd expression could contribute to variation in ET from FNR to SIR and complementation of *E. coli* EW11 growth^25^. To test this idea, we fused each chim-Fd to red fluorescent protein (RFP) at their C-terminus using a (GSS)_4_ linker and examined protein expression. This linker was chosen because we previously showed that it can be used to fuse RFP to the C-terminus of ml-Fd without disturbing cellular ET^35^. With each Fd, whole cell fluorescence measurements revealed a higher signal in the presence of aTc compared with uninduced cultures (Figure 4B). ml-Fd1-RFP presented the largest aTc-dependent fluorescence. Most of the chim-Fds presented signals that were ≥50% of that observed with the parental Fds, with the exception of chim-Fd-1 and chim-Fd-7, which presented significantly lower signals. In some cases, the protein expression did not correlate with growth complementation. For example, chim-Fd7 presented the weakest growth complementation, but did not present the lowest expression.

### Chimera midpoint reduction potentials

Electrochemical changes arising from recombination could also contribute to the variation in *E. coli* EW11 complementation, since growth requires ET between partner proteins with defined midpoint potentials, zmFNR (E° = −337 mV) to zm-SIR (E°= −285 mV)^36,37^. To determine how the chim-Fd midpoint reduction potentials relate to those of the donor and acceptor proteins, we performed protein thin-film, square wave voltammetry on each chim-Fd. Surprisingly, all of the chimeras presented E° outside of the range of the parental proteins (−344 mV to - 336 mV) and all but two were more negative than the parents (Figure 5A; S4; S5). Several chimeras presented midpoint potentials lower than the most negative parent (ml-Fd1). The two chimeras presenting positively shifted E° from the parents had smaller shifts than those that were negatively shifted. These results can be contrasted with the T_m_ trends, which yielded values bounded by the parental values (Figure 5B). While chim-Fds presented E° and T_m_ values that both differed from the parental Fds, these properties did not display a significant correlation (r = −0.55, p > 0.1) (Figure S6).

**Figure 5.**
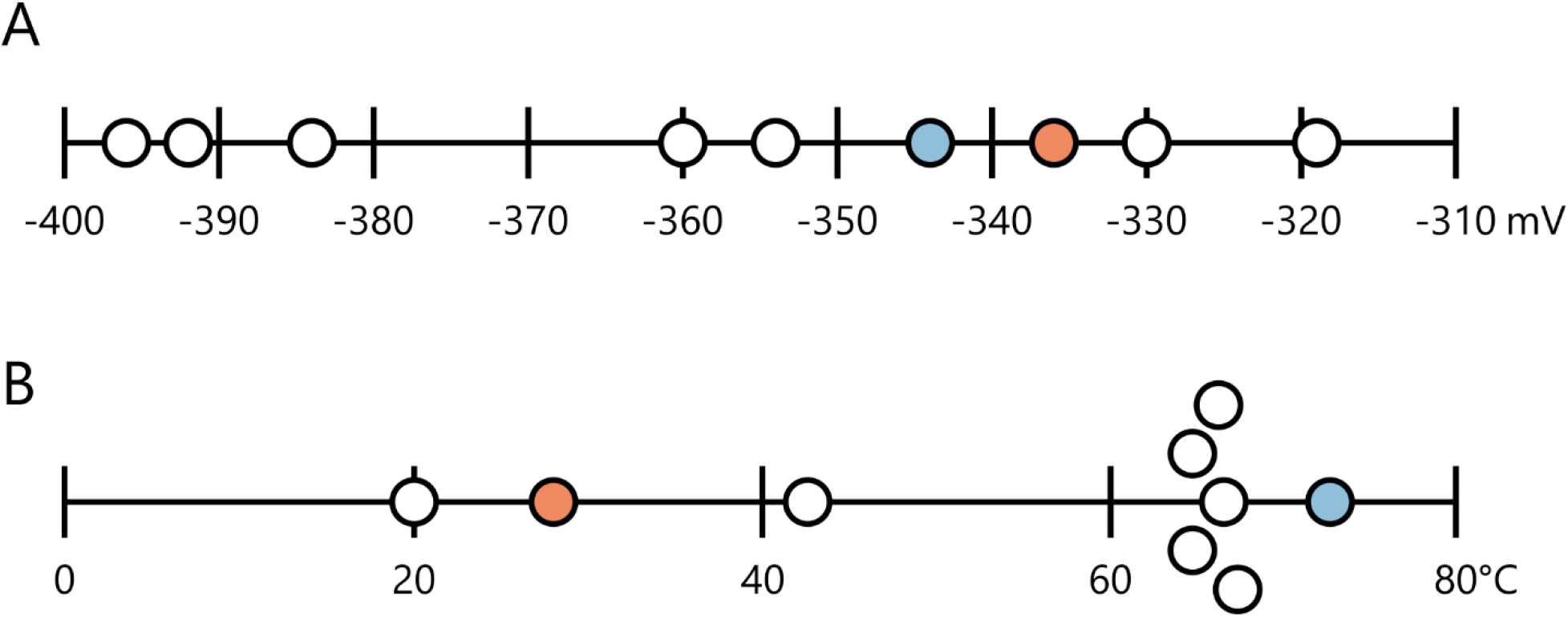
Midpoint potential and thermostability differ in inheritance patterns. (**A**) The E° of purified ml-Fd1 (blue), pssm2-Fd (orange) are compared with the chim-Fds (white). E° were measured using square-wave thin-film protein voltammetry with concentrated protein samples (300 μM) at pH 7 and 23.5°C. (**B**) The T_m_ of chim-Fds (white) reveals values that are largely intermediate to ml-Fd1 (blue) and pssm2-Fd (orange). All thermostability measurements were performed by monitoring ellipticity (427 nm) of samples containing 50 μM protein in TED buffer at a scan rate of 1 C°/min.

### Biophysical comparisons

To better understand the effects of recombination on protein thermostability, we compared the T_m_ values of each Fd-chim with a range of physicochemical properties. A comparison of T_m_ with the calculated isoionic point (pI) revealed a significant correlation (r = 0.87, p < 0.005) (Figure 6A), as well as between pI and the absolute counts of charged amino acids in each chim-Fd (Figures S7A-C). These findings support the idea that electrostatics play a role in controlling the stabilities of Fds^27,38,39^. A comparison of T_m_ with mutational distance from ml-Fd1 (Figure 6B) yielded an even stronger correlation (r = 0.94, p < 0.005). Since some mutations disrupt structure by breaking residue-residue contacts critical to folding and function, we also evaluated whether the number of residue-residue contacts broken (C_broken_) through recombination correlates with Tm^9,13,28,40,41^. This analysis showed that chim-Fds tolerate up to 20 broken residue-residue contacts broken before losses in T_m_ are observed (Figure 6C). When this latter analysis was performed by calculating contacts broken of all chim-Fds relative to one of the parental structures, similar trends were observed (Figures S8A-B). We also evaluated whether the E° of the chim-Fds correlate with the same parameters. In all cases, the trends revealed weaker correlations (Figures 6D-6F; S7D-F; S8C-D), none of which were significant.

**Figure 6.**
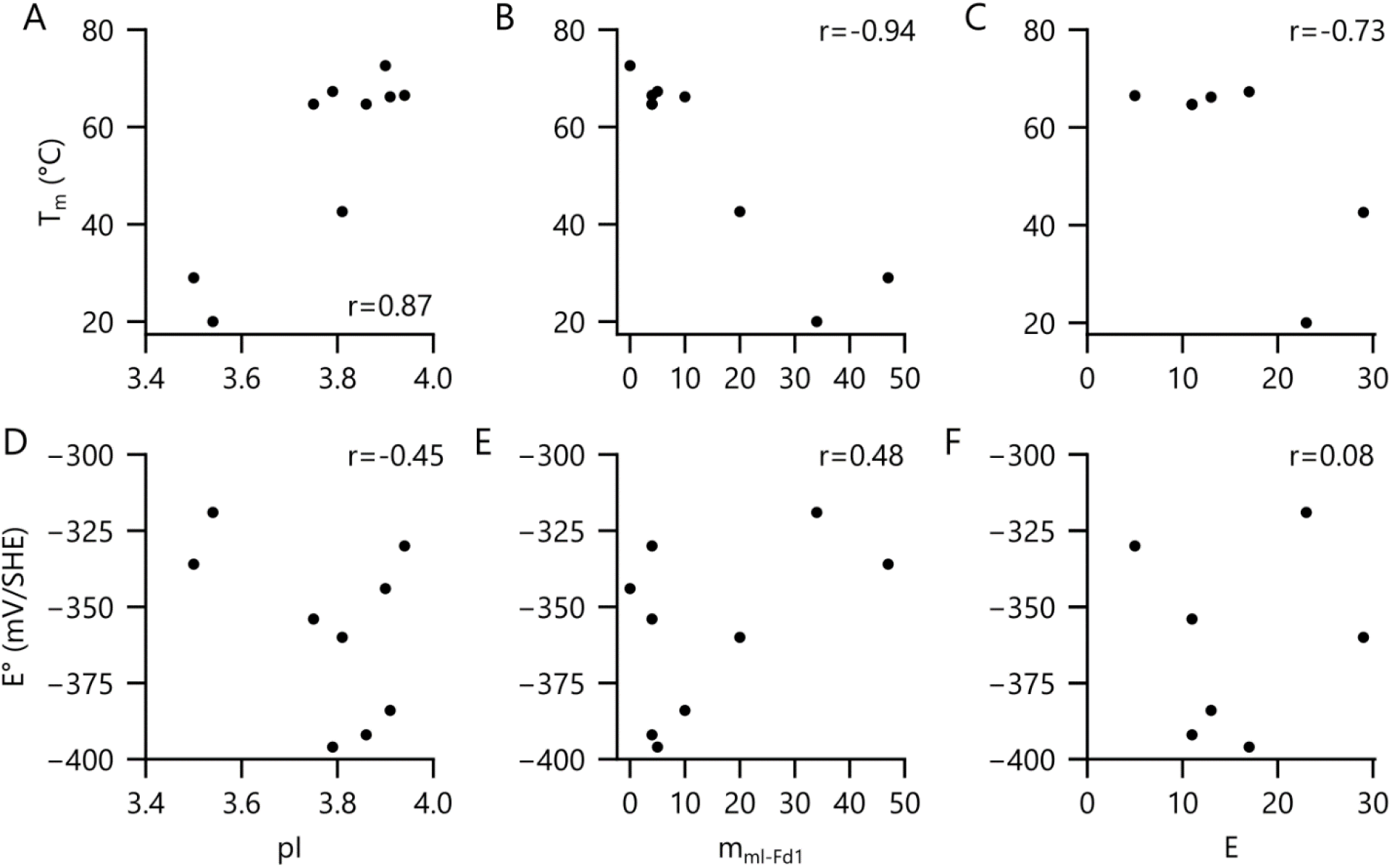
Biophysical correlations. The T_m_ values of Fds are compared to (**A**) calculated pI values, (**B**) mutational distance from the more thermostable ml-Fd1, m_ml-fd1_, and (**C**) residue-residue contacts broken by recombination, C_broken_. The E° of each Fd is compared to (**D**) pI, (**E**) m_ml-fd1_, and (**F**) C_broken_. Pearson correlations are shown in each panel. Significant correlations were observed between T_m_ and pI (p<0.005) and T_m_ and m_ml-Fd1_ (p<0.0005). All other trends presented p values >0.05.

### Implications for oxidoreductase design

Our results provide evidence that chim-Fds created by recombining distantly-related parents (53% identity) display a high tolerance to amino acid substitutions. All of the chim-Fds characterized herein retained the ability to coordinate an iron-sulfur cluster, and they all supported ET from FNR to SIR in a cellular assay, even though they have a wide range of amino acid substitutions (4 to 34) and contacts disrupted by recombination (5 to 29). These findings are similar to those obtained in a study examining the folding and function of chimeric cytochromes P450, which found that a vast majority of the chimeras having ≤35 amino acid substitutions and ≤35 disrupted residue-residue contacts broken retained the ability to fold, bind a metallocofactor, and perform catalysis^13^. In contrast, studies examining the function of lactamase chimeras created by recombining parents with 40% identity found that only a subset of the chimeras with ≤35 disrupted residue-residue contacts retained function^41,42^. With lactamases, a thermostabilizing mutation could increase protein tolerance to recombination^42^. Additionally, chimeric cellulases created by recombining distantly related parents (64% identity) exhibited a greater sensitivity to disrupted residue-residue contacts^40^. Cellulase chimeras having ≥15 broken residue-residue contacts were largely inactive. Taken together, these findings suggest that metal-dependent oxidoreductases may tolerate a higher number of broken residue-residue contacts compared with enzymes like lactamases and cellulases.

With homologous recombination, melting temperatures of the resulting chimeras have been found to correlate with the polypeptide inheritance from parental proteins. When three cytochromes P450 were recombined with distinct melting temperatures, the distribution of chimera thermostabilities could be predicted from the additive contributions of the sequence fragments inherited^12,20^. A vast majority of the chimeras presented melting temperatures that were intermediate to those of the parents, and chimeras only rarely exhibited higher or lower melting temperature values. Our results with chimeric Fds are consistent with these observations. All of the chim-Fds exhibit melting temperatures that are intermediate to the parental proteins recombined. In contrast, the electrochemical properties of the chimeric Fds are all outside the range of the parental proteins. This trend is distinct from that observed in prior Fd recombination studies, which found that the E° of chimeras were bounded by the values of the parental proteins^15–17^. The underlying cause of the contrasting trend observed herein not known. The two Fd chimeras presenting the largest midpoint reduction potential shifts arise from sequence changes on the β-strand most distal to the [2Fe-2S] cluster, although each chimera arises from mutations on opposite ends of the strand. Additionally, chim-Fd5, which also presented a low E°, has sequence changes adjacent to a cluster-ligating cysteine.

Our results calibrate the effects of recombining distantly-related Fds, which will be useful in the future for guiding the design of larger chimeric libraries. The high tolerance of Fd structure and cellular ET to recombination suggests that libraries created by recombining distantly-related family members will contain a high fraction of folded variants that retain the ability to coordinate an iron-sulfur cluster. By selecting large libraries of chim-Fds for variants that support ET between zm-FNR and zm-SIR, one could rapidly identify variants that are most efficient in this cellular pathway, since growth depends on Fd-mediated ET. By varying FNR and SIR partners in this selection, which may favor ET with distinct chimeras, selections could provide insight into the ways that Fd ET efficiencies in cells depend upon the structure of their partner proteins. By analyzing how the physical properties of chim-Fds vary with primary structure and cellular function, it may be possible to begin developing design rules that guide the creation of Fds with user specified thermostabilities, midpoint potentials, and cellular ET.

## METHODS AND MATERIALS

### Materials

Tris Base was from Fisher Scientific, N-cyclohexyl-3-aminopropanesulfonic (CAPS) was from Acros Organics, 2-(N-Morpholino)ethanesulfonic acid (MES) and N-[Tris(hydroxymethyl)methyl]-3-aminopropanesulfonic acid (TAPS) was from Fluka Biochemika. Isopropyl β-D-1-thiogalactopyranoside (IPTG), dithiothreitol (DTT), kanamycin, chloramphenicol, and streptomycin were from Research Product International. 3-Morpholinopropane-1-sulfonic acid (MOPS), N-Cyclohexyl-2-aminoethanesulfonic acid (CHES), and all other chemicals were purchased from Sigma-Aldrich. *E. coli* EW11 was a gift from Pam Silver (Harvard University)^34^, *E. coli* XL1-Blue was from Agilent, and *E. coli* Rosetta™(DE3) was from Novagen.

### Vector Design

All plasmids used for cellular measurements are listed in Table S1, while those used to overexpress proteins for purification are in Table S2. Genes were synthesized by Integrated DNA technologies as G-blocks. Plasmids were constructed by ligating PCR products amplified with Q5^®^ High-Fidelity DNA polymerase (New England Biolabs) using Golden Gate DNA assembly^43^. Ribosome binding sites were designed using the RBS calculator^44^. All Fds had the same translation initiation site, except for pssm2-Fd. All plasmids were sequence verified.

### Calculations

Alignments were made using MUSCLE and visualized using JalView^45,46^. pI values were estimated using ExPASy^47^. Residue-residue contacts broken by recombination was calculated using SCHEMA^9^. Unless noted otherwise, disruption of all chimeras was calculated relative to either designated parent or parent most similar to each chimera and it is reported as a disruption score (C_broken_). Pairwise amino acids were only counted if residues, including both sidechain and main chain atoms, were within 4.5 Å of each other and recombination caused a change in sequence relative to parents. Hydrogen bonds and salt bridges were identified using PyMol^48^.

### Protein purification

*E. coli* Rosetta™(DE3) transformed with pET28b-derived vectors containing Fds were grown at 37°C in lysogeny broth (LB) containing 50 μg/mL kanamycin to exponential phase, induced at midlog phase using 50 μM IPTG, and grown overnight at 37°C while shaking at 250 rpm. Cells harvested by centrifugation (4,000 g) were resuspended in lysis buffer, which contained 10 mM Tris pH 8, 5 mM dithiothreitol (DTT), 10 mg/L DNase I, and 0.5 mg/mL lysozyme. After freezing at −80°C, cells were thawed and mixed with cOmplete Mini, EDTA-Free protease inhibitor (Sigma-Aldrich) at a ratio of one tablet per 100 mL lysate. Clarified lysate harvested by centrifugation was diluted 3-fold with TED buffer (25 mM Tris pH 8, 1 mM EDTA, 1 mM DTT) and loaded onto a DE52 anion exchange column (Whatman). The column was washed with TED containing 200 mM NaCl, and the Fd was eluted using sequential isocratic washes with TED containing 250 and 300 mM NaCl. Fractions appearing brown were mixed, diluted with TED, and loaded onto HiTrap Q XL column (GE Healthcare) using an AKTA Start FPLC system (GE Healthcare). After washing the column with TED, the protein was eluted using a linear gradient (0 to 375 mM NaCl in TED) followed by an isocratic wash (500 mM NaCl in TED). Brown fractions were pooled and then purified using a HiLoad 16/600 Superdex 75 (GE Healthcare) size exclusion column containing TED. SDS-PAGE was performed to analyze purity at each step using NuPage 12% Bis-Tris Gels (Invitrogen). Samples appearing homogeneous were pooled and concentrated using an Amicon Ultra 10 K MWCO spin column (EMD Millipore) and flash frozen with liquid nitrogen.

### Spectroscopy

To obtain buffer matched controls, Fds were dialyzed into TED prior to all measurements. Absorbance spectra and ellipticity were acquired using a J-815 spectropolarimeter (Jasco, Inc) using quartz cuvettes with a 1 cm path length. Scans were conducted using a 1 nm bandwidth, a 0.5 nm data pitch, and a 200 nm/min scan rate at 20°C. To assess protein stability, a cuvette containing 50 μM Fd was heated from 5 to 95°C at a rate of 1°C/minute while monitoring ellipticity and absorbance. All spectra represent buffer corrected data.

### Electrochemistry

Electrochemical measurements were performed anaerobically using a three-electrode system. A Ag/AgCl/1M KCl electrode (CH Instruments) was used as the reference electrode, and a platinum wire was used as the counter electrode. An edge-plane pyrolytic graphite electrode was used as the working electrode to perform protein film electrochemistry. Prior to adding Fd, this electrode was treated with 100 mM neomycin trisulfate (Sigma-Aldrich) to improve the electrochemical signal^49^. An aliquot (3μL) of Fd (~300 uM) was applied directly to the electrode surface following neomycin treatment, and the protein was allowed to adhere to the surface for 1 min at 23°C. The electrode was then placed in a glass vial containing a pH 7 buffer solution (5 mM acetate, MES, MOPS, TAPS, CHES, CAPS) containing 100 mM NaCl at 23.5 °C. Square wave voltammograms were collected at 10 Hz frequency, and electrochemical signals were analyzed using Qsoas open software. Similar results were obtained when experiments were performed using pssm2-Fd from different purifications. A CH Instruments potentiostat and CHI660E electrochemical analyzer were used for all measurements. All data is reported relative to Standard Hydrogen Electrode (SHE), taking into account the potential difference between SHE and Ag/AgCl/1M KCl, which is 0.222 V.

### Growth assay

*E. coli* EW11 cells were transformed with two plasmids using electroporation, one constitutively expressing the electron donor and acceptor pair (zm-FNR and zm-SIR) and the other expressing a native Fd or a chimera as previously described^24,34,35,50^. To select for the Fd and partner plasmids, all growth steps included chloramphenicol (34 μg/mL) and streptomycin (100 μg/mL). Starter cultures were inoculated using single colonies. These starter cultures were grown in deep-well 96-well plates for 18 h at 37 °C in 1 mL of a non-selective modified M9 medium (M9c) as previously described^25^. Starter cultures that had been grown to stationary phase were centrifuged at 4 °C for 10 min at 3500 xg and resuspended in 1 mL of a selective modified M9 medium (M9sa), which is identical to M9c but lacks cysteine and methionine. Starter cultures were then diluted 1:100 into M9sa in Nunc™ Edge 2.0 96-well plates (Thermo Fisher). Cells were grown in the presence of the indicated amount of aTc in a Spark plate reader (Tecan) at 37 °C with shaking at 90 rpm at an amplitude of 3 mm in double-orbital mode. Optical density at 600 nm (OD_600_) was measured for 48 h.

### Cellular fluorescence

These measurements were performed like the growth assay except cultures were grown in M9c throughout. After 48 h, OD_600_ and emission (λ_ex_ = 560 nm; λ_em_ = 650 nm) were measured. All values shown represent fluorescence normalized to OD_600_.

### Statistics

Error bars represent standard deviation calculated from three or more biological replicates. Independent, two-tailed t-tests were used to compare differences between all relevant samples with α=0.05. All correlations shown are Pearson correlations calculated with NumPy^51^.

## SUPPORTING INFORMATION

The Supporting Information includes:

**Table S1:** List of vectors used for cellular assay.

**Table S2:** List of vectors used for overexpression and purification.

**Figure S1:** Structural models of chimeric Fds.

**Figure S2:** Absorbance spectra of chimeric Fds.

**Figure S3:** Iron-sulfur cluster content of chimeric Fds.

**Figure S4:** Buffer-subtracted square wave voltammetry of chimeric Fds.

**Figure S5:** Baseline square wave voltammetry of chimeric Fds.

**Figure S6:** Comparison of chimera thermostability and midpoint potential.

**Figure S7:** Comparisons of chimera amino acid composition, thermostability, and midpoint potential.

**Figure S8:** Correlations between residue-residue contacts broken by recombination, thermostability, and midpoint potential.

## ACKNOWLEDGMENTS

This project was supported by DOE grant DE-SC0014462 (to J.J.S. and G.N.B.), NASA NAI grant 80NSSC18M0093 (to G.N.B and J.J.S.), NSF grant 1843556 (to G.N.B., J.J.S., and R.V.). I.J.C. and J.T.A. were supported by a Lodieska Stockbridge Vaughn Fellowship. C.P.T. is a participant in the NSF Research Traineeship 1828869 (to J.J.S., G.N.B., and R.V.).

## Notes

### Competing Interest Statement

The authors have declared no competing interest.

