## Supporting Information for "Recombination of 2Fe-2S ferredoxins reveals differences in the inheritance of thermostability and midpoint potential"

### List of contents

1. **Table S1**
2. **Table S2**
3. **Figure S1**
4. **Figure S2**
5. **Figure S3**
6. **Figure S4**
7. **Figure S5**
8. **Figure S6**
9. **Figure S7**
10. **Figure S8**

**Table S1. Assay vectors used.** For each vector, the name, antibiotic marker, origin, proteins expressed, and Addgene IDs are noted. n/s not submitted.

| Name | Description | Addgene |
|---|---|---|
| pSAC01 | $Spec^R$, p15a vector that expresses *zm*-FNR & zm-SIR | 131826 |
| pFd007 | $Cam^R$, ColE1 vector with aTc inducible *ml*-Fd1 | 131828 |
| pFd022.R2 | $Cam^R$, ColE1 vector with aTc inducible pssm2-Fd | 137967 |
| pDC2 | $Cam^R$, ColE1 vector with aTc inducible *chim-Fd1* | n/s |
| pDC6 | $Cam^R$, ColE1 vector with aTc inducible chim-Fd2 | n/s |
| pDC19 | $Cam^R$, ColE1 vector with aTc inducible chim-Fd3 | n/s |
| pDC23 | $Cam^R$, ColE1 vector with aTc inducible chim-Fd4 | n/s |
| pDC22 | $Cam^R$, ColE1 vector with aTc inducible chim-Fd5 | n/s |
| pDC28 | $Cam^R$, ColE1 vector with aTc inducible chim-Fd6 | n/s |
| pDC36 | $Cam^R$, ColE1 vector with aTc inducible chim-Fd7 | n/s |
| pFd007.RFP | $Cam^R$, ColE1 vector with aTc inducible *ml*-Fd1-RFP | n/s |
| pFd022.R2.RFP | $Cam^R$, ColE1 vector with aTc inducible *pssm2-Fd*-RFP | 137971 |
| pDC2_L12_RFP | $Cam^R$, ColE1 vector with aTc inducible chim-Fd1-RFP | n/s |
| pDC6_L12_RFP | $Cam^R$, ColE1 vector with aTc inducible chim-Fd2-RFP | n/s |
| pDC19_L12_RFP | $Cam^R$, ColE1 vector with aTc inducible chim-Fd3-RFP | n/s |
| pDC23_L12_RFP | $Cam^R$, ColE1 vector with aTc inducible chim-Fd4-RFP | n/s |
| pDC22_L12_RFP | $Cam^R$, ColE1 vector with aTc inducible chim-Fd5-RFP | n/s |
| pDC28_L12_RFP | $Cam^R$, ColE1 vector with aTc inducible chim-Fd6-RFP | n/s |
| pDC36_L12_RFP | $Cam^R$, ColE1 vector with aTc inducible chim-Fd7-RFP | n/s |

**Table S2. Vectors used for overexpression and purification.** For each vector, the name, antibiotic marker, origin, proteins expressed, and Addgene IDs are noted. n/s not submitted.

| Name | Description | Addgene |
|---|---|---|
| pJTA007 | $Kan^R$, ColE1 pET28b derived vector with T7-lac inducible *ml*-Fd1 | 132328 |
| pJTA022 | $Kan^R$, ColE1 pET28b derived vector with T7-lac inducible *pssm2*-Fd | 137975 |
| pEX.DC2 | $Kan^R$, ColE1 pET28b derived vector with T7-lac inducible chim-Fd1 | n/s |
| pEX.DC6 | $Kan^R$, ColE1 pET28b derived vector with T7-lac inducible chim-Fd2 | n/s |
| pEX.DC19 | $Kan^R$, ColE1 pET28b derived vector with T7-lac inducible chim-Fd3 | n/s |
| pEX.DC23 | $Kan^R$, ColE1 pET28b derived vector with T7-lac inducible chim-Fd4 | n/s |
| pEX.DC22 | $Kan^R$, ColE1 pET28b derived vector with T7-lac inducible chim-Fd5 | n/s |
| pEX.DC28 | $Kan^R$, ColE1 pET28b derived vector with T7-lac inducible chim-Fd6 | n/s |
| pEX.DC36 | $Kan^R$, ColE1 pET28b derived vector with T7-lac inducible chim-Fd7 | n/s |

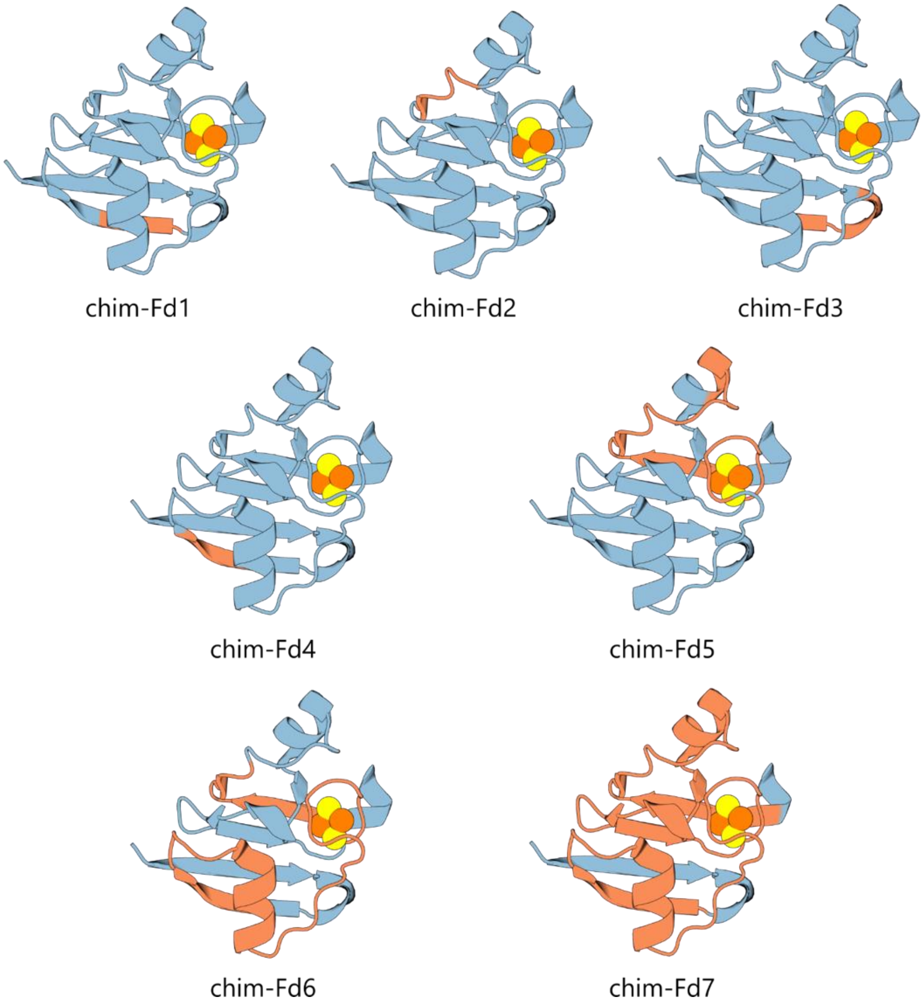


**Figure S1. Structural models of chimeric Fds.** The inheritance of ml-Fd1 structure (blue) and pssm2-Fd show sites of recombination used to generate chim-Fds.

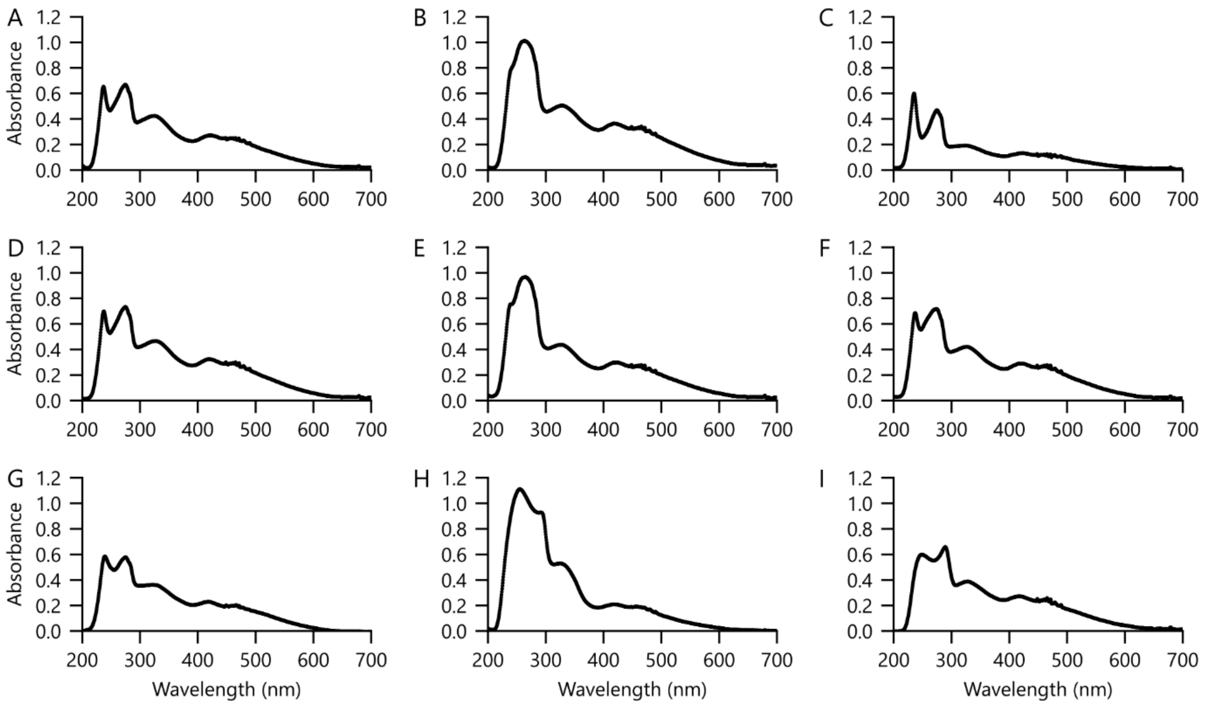


**Figure S2. Absorbance spectra of Fds.** The absorbance spectra of (**A**) ml-Fd1, (**B**) chim-Fd1, (**C**) chim-Fd2, (**D**) chim-Fd3, (**E**) chim-Fd4, (**F**) chim-Fd5, (**G**) chim-Fd6, (**H**) chim-Fd7, and (**I**) pssm2-Fd. Measurements were performed in TED buffer at 23°C, and were accumulated 3 times before averaging .

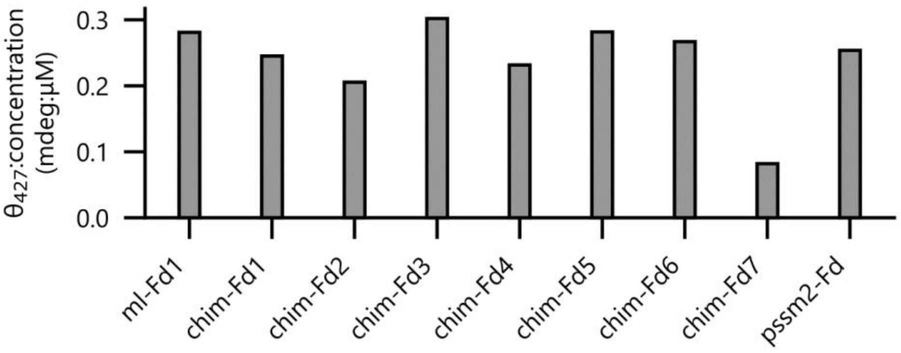


**Figure S3. Effect of recombination on iron-sulfur cluster incorporation.** The ratio of ellipticity at 427 nm to concentration for each chimeric Fd is compared with the parental proteins ml-Fd1 and pssm2-Fd.

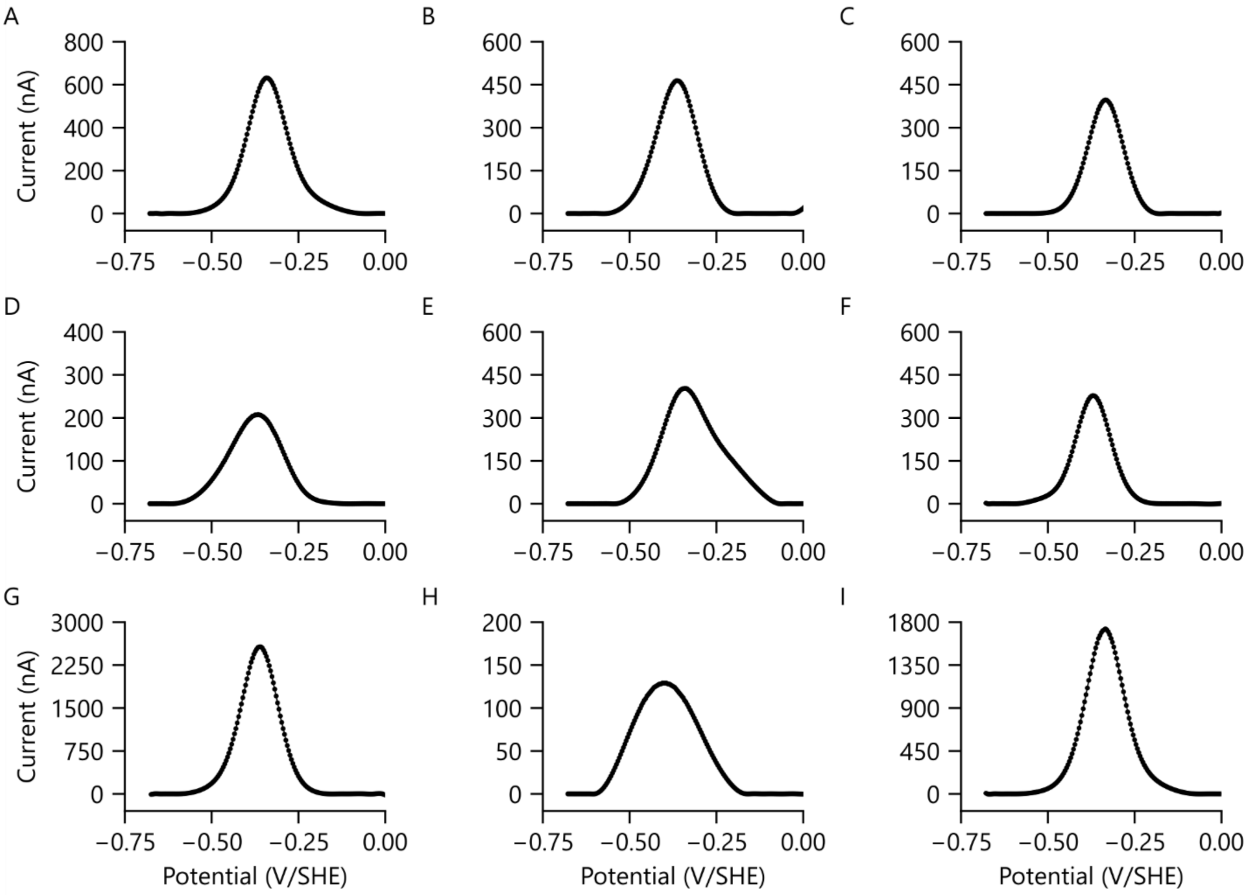


**Figure S4.**The square-wave thin-film protein voltammetry of purified (**A**) ml-Fd1, (**B**) chim-Fd1, (**C**) chim-Fd2, (**D**) chim-Fd3, (**E**) chim-Fd4, (**F**) chim-Fd5, (**G**) chim-Fd6, (**H**) chim-Fd7, and (**I**) pssm2-Fd after buffer subtraction. Voltammetry was performed with concentrated protein samples (300 μM) at pH 7 and 23.5°C.

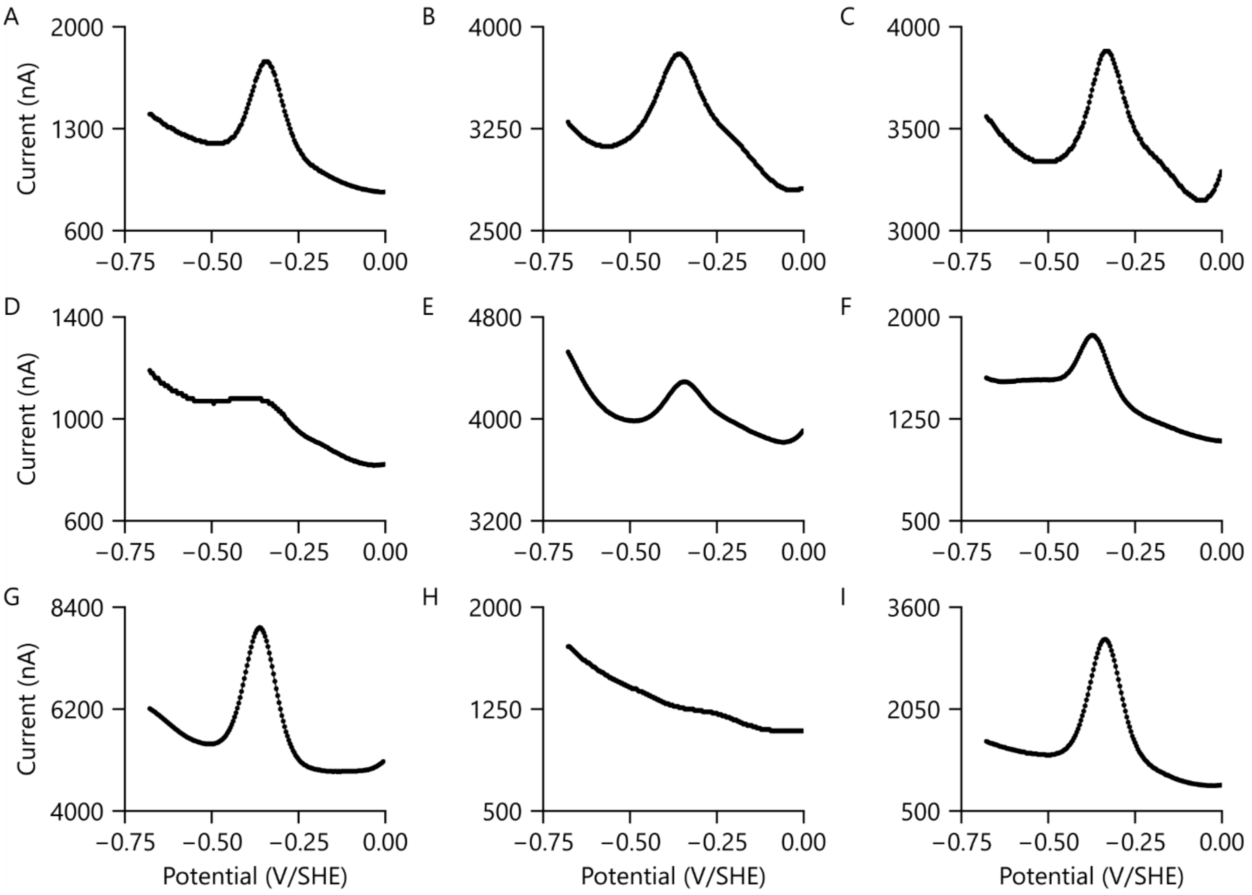


**Figure S5.**The baseline square-wave thin-film protein voltammetry of purified (**A**) ml-Fd1, (**B**) chim-Fd1, (**C**) chim-Fd2, (**D**) chim-Fd3, (**E**) chim-Fd4, (**F**) chim-Fd5, (**G**) chim-Fd6, (**H**) chim-Fd7, and (**I**) pssm2-Fd. Voltammetry was performed with concentrated protein samples (300 µM) at pH 7 and 23.5°C.

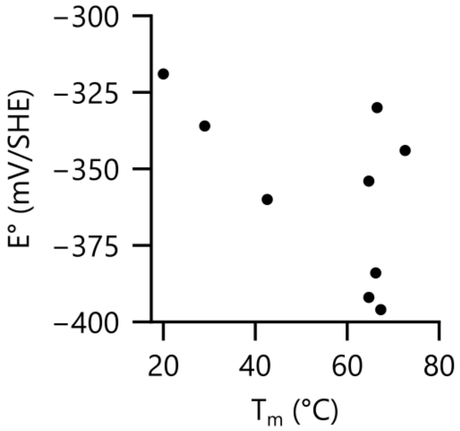


**Figure S6. Comparison of thermostability and midpoint potential.** A comparison of Fd melting temperatures ($T_m$) are E° (mv/SHE) does not present a significant correlation (Pearson correlation = -0.55).

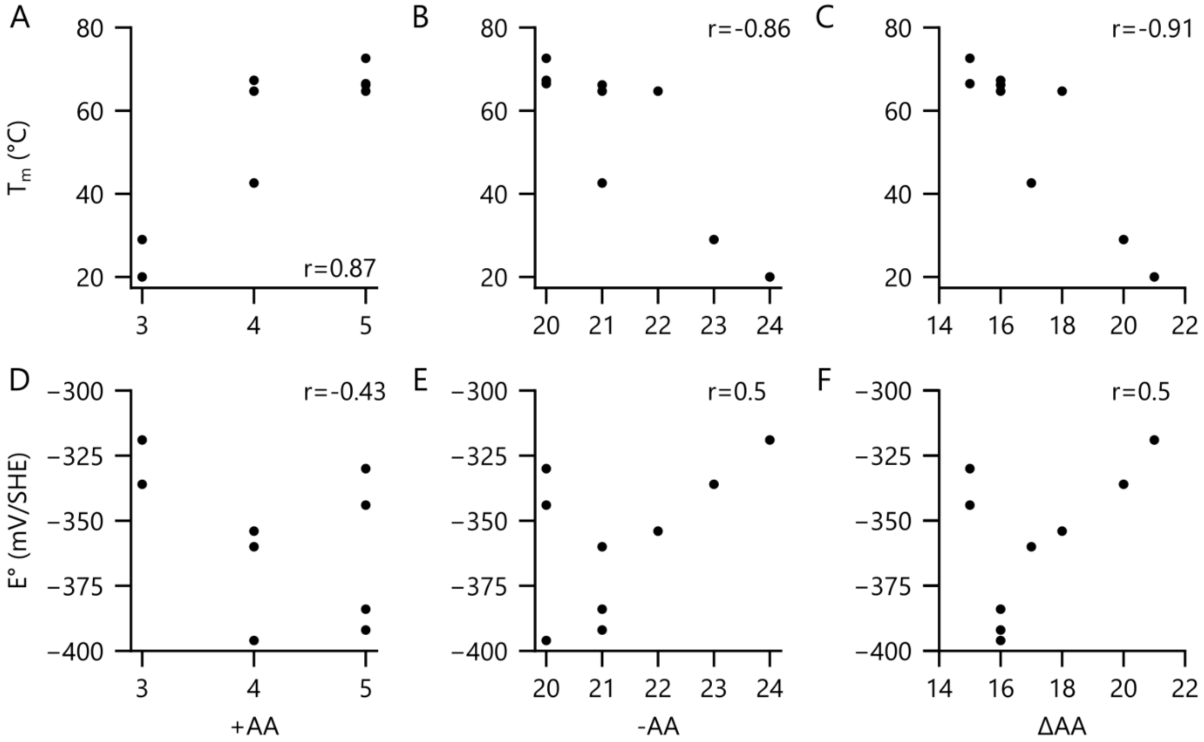


**Figure S7. Correlations of $T_m$ and E° with charged amino acid content.** Fd melting temperatures are compared to (**A**) counts of positively charged amino acids (+AA), (**B**) counts of negatively charged amino acids (-AA), and (**C**) the count of negative amino acids subtracted by the count of positive amino acids (ΔAA). Fd midpoint reduction potentials are compared to (**D**) +AA, (**E**) -AA, and (**F**) ΔAA. Pearson correlations are listed on each chart. Significant correlations were observed between $T_m$ and +AA ($p<0.005$), $T_m$ and -AA($p<0.005$), and $T_m$ and ΔAA ($p<0.005$). All other trends presented p values $>0.05$.

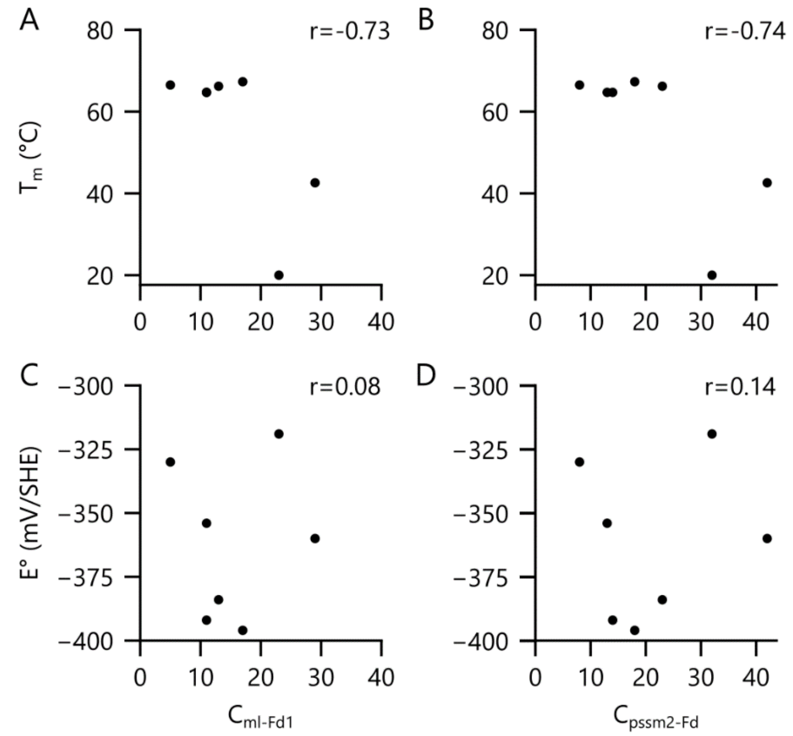


**Figure S8. Correlations of $T_m$ and E° with residue-residue contacts broken by recombination.** Fd melting temperatures are compared with broken residue-residue contacts calculated using (**A**) the ml-Fd1 structure, $C_{ml\text{-}Fd1}$, and (**B**) the pssm2-Fd structure, $C_{pssm2\text{-}Fd}$. Parental and chimeric Fd midpoint potentials are also compared to (**C**) $C_{ml\text{-}Fd1}$ and (**D**) $C_{pssm2\text{-}Fd}$. Pearson correlations are listed on each chart. All trends presented p values >0.05.
